# Relationships between the hard and soft dimensions of the nose in *Pan troglodytes* and *Homo sapiens* reveal the nasal protrusions of Plio-Pleistocene hominids

**DOI:** 10.1101/2021.10.18.464897

**Authors:** Ryan M. Campbell, Gabriel Vinas, Maciej Henneberg

## Abstract

By identifying similarity in bone and soft tissue covariation patterns in hominids, it is possible to produce facial approximation methods that are compatible with more than one species of primate. In this study, we conducted an interspecific comparison of the nasomaxillary region in chimpanzees and modern humans with the aim of producing a method for predicting the nasal protrusions of ancient Plio-Pleistocene hominids. We addressed this aim by first collecting and performing regression analyses of linear and angular measurements of nasal cavity length and inclination in modern humans (*Homo* sapiens; n = 72) and chimpanzees (*Pan troglodytes; n* = 19), and then by performing a set of out-of-group tests. The first test was performed on two subjects that belonged to the same genus as the training sample, i.e., *Homo* (*n* = 1) and *Pan* (*n* = 1), and the second test, which functioned as an interspecies compatibility test, was performed on *Pan paniscus* (*n* = 1), *Gorilla gorilla* (*n* = 3), *Pongo pygmaeus* (*n* = 1), *Pongo abelli* (*n* = 1), *Symphalangus syndactylus* (*n* = 3), and *Papio hamadryas* (*n* = 3). We identified statistically significant correlations in both humans and chimpanzees with slopes that displayed homogeneity of covariation. Joint prediction formulae were found to be compatible with humans and chimpanzees as well as all other African great apes, i.e., bonobos and gorillas. The main conclusion that can be drawn from this study is that regression models for approximating nasal projection are homogenous among humans and African apes and can thus be reasonably extended to ancestors leading to these clades.

## Introduction

The process of producing faces from dry skulls is known as facial approximation. Since the purpose of this procedure is to estimate a subject’s premortem anatomy as closely as possible, each facial feature requires a robust scientific method. While each feature of the face is important in its own right, accurate prediction of nasal protrusion is critical because the nose is in the centre of the face. Approximation error related to nasal protrusion can thus significantly change the facial appearance of the subject in question. For this reason, the tip of the nose (pronasale) is a prominent landmark in the forensic facial approximation literature (Gerasimov, 1955, 1971; Lee et al., 2014; Rynn et al., 2010; Stephan et al., 2003; Wilkinson, 2004).

The pronasale landmark is equally important in facial approximations of extinct Plio-Pleistocene hominids; in this paper hominids means all members of the Hominidae, which is comprised of the African apes, humans, and all ancestors leading to these clades. It has been stated that primate comparative anatomy, which is the study of similarities and differences in structures of different species, is critical to the practice of ancient hominid facial approximation. However, despite numerous facial approximations of extinct hominids presented in scientific textbooks and museum displays, interspecific variation in soft tissue nasal form between humans and chimpanzees has received little research interest. Although some overlap between human and chimpanzee noses is recognized to exist, why modern humans possess a particularly unique projecting, external nose is essentially a mystery. In contrast to human noses, the noses of chimpanzees, and of other great apes (bonobos, gorillas, and orangutans), are relatively flat. An investigation into the morphological differences between extant hominids may result in more accurate facial approximation methods, which are needed to reduce the excessive variability recognized in facial approximations of the same individual (Anderson, 2011; Campbell et al., 2021b).

In the facial approximation literature, eight methods for approximating the nasal profile in modern humans have been published (George, 1987; Gerasimov, 1955, 1971; Krogman, 1962; Macho, 1986; Prokopec and Ubelaker, 2002; Rynn et al., 2010; Stephan et al., 2003). Studies testing these methods (Rynn and Wilkinson, 2006; Stephan et al., 2003) have consistently reported that the method by George (1987) appears to be the most useful. It consists of calculating a percentage (60.5% for males and 56% for females) of a distance from nasion to the inferior nasal spine to establish a chord at subnasale parallel to the Frankfurt horizontal plane. Similarly, orthodontists and maxillofacial surgeons, who share an interest in nasal morphology, have identified numerous correlations between the soft and hard nasal tissues (Jankowska et al., 2021). These correlations were first explored in Stephan et al. (2003) and then in Rynn et al. (2010). Both of these studies have produced regression equations for approximating nasal morphology from dry skulls, although the method by Stephan et al. (2003) has been shown to underestimate nasal protrusion (Rynn and Wilkinson, 2006). Regardless of the validity and reliability of these methods, there is still no evidence supporting their use on fossil hominids.

Most evolutionary studies of the nasal region have focused on modern humans (Bastir and Rosas, 2013; Fukase et al., 2016; Maddux et al., 2016) and Neanderthals/Neandertals (de Azevedo et al., 2017; Holton and Franciscus, 2008; Laitman et al., 1996; Márquez et al., 2014; Schwartz et al., 2008). Conversely, very little attention has been paid to the soft tissue of great ape noses. While we acknowledge that chimpanzee noses have received some research interest (Losken et al., 1994; Samarat, 2016; Samarat et al., 2013), with the exception of one study (Hofer, 1972), gorilla and orangutan studies are practically non-existent. It is not clear why the nasal soft tissues of great apes are so understudied. However, it is likely a direct result of the status of these animals as endangered species and the current difficulties involved in obtaining this kind of material (pers. observation).

It has been said that among all the transformations in craniofacial morphology recognized to have occurred during the proposed—although contested (see Kimbel and Villmoare, 2016)—transition from *Australopithecus* to *Homo*, nasal morphology is a major one (Franciscus and Trinkaus, 1988a). In particular, the nose of modern humans is distinguished from great apes by possessing an external part that protrudes past the piriform aperture. The evolutionary reasons for this feature have been the subject of continuing scientific debate. Some studies argue that the external nose was derived in the genus *Homo* because of adaptation to climate (Noback et al., 2011; Weiner, 1954; Wolpoff, 1968; Zaidi et al., 2017). According to one hypothesis (Franciscus and Trinkaus, 1988a), the nose arose out of the face through expansion of the nasal bones relative to that of the piriform aperture. This is said to have occurred as Pleistocene hominids, such as *Homo erectus*, shifted to increasingly prolonged bouts of physical activity in arid environments, resulting in selective pressures on adaptive respiratory function (Franciscus and Trinkaus, 1988a). In contrast, and more recently, Nishimura et al. (2016) showed that the external protruding nose in modern humans has little effect on improving air conditioning. They concluded that the unique nasal anatomy in *Homo* was likely formed passively by facial reorganization and not from adaptation to climate. It is important to note that neither of these explanations provide the empirical data needed in forensic facial approximations of Plio-Pleistocene hominids. In addition, the question of whether the protrusion growth of the nose was constant or punctuated is entirely unanswered. Obtaining this knowledge is crucial to inform practitioners of facial approximation of how to model the nasal anatomy of their subjects and produce scientifically accurate approximations of hominids. Thus, further scientific research is needed to compare the nasal region among extant hominids.

The aims of this study were to explore this matter further. We aimed to (1) compare pronasale position in modern humans and chimpanzees, and (2) to produce prediction formulae for approximating the nasal protrusions of ancient Plio-Pleistocene hominids. We addressed these aims twofold: Firstly, by collecting and performing regression analyses of linear and angular measurements of nasal cavity length and inclination in modern humans (*Homo* sapiens; n = 72) and chimpanzees (*Pan troglodytes*; *n* = 19); and secondly, by performing a set of out-of-group tests. The first test was performed on two subjects that belonged to the same genus as the training sample, i.e., *Homo* (*n* = 1) and *Pan* (*n* = 1), and the second test, which functioned as an interspecies compatibility test, was performed on *Pan paniscus* (*n* = 1), *Gorilla gorilla* (*n* = 3), *Pongo pygmaeus* (*n* = 1), *Pongo abelli* (*n* = 1), *Symphalangus syndactylus* (*n* = 3), and *Papio hamadryas* (*n* = 3). We hypothesize that soft tissue approximation models, such as those for nasal protrusion, are homogenous among extant hominids and can thus be reasonably extended to all ancestors leading to these clades. To illustrate this hypothesis, we approximated the nasal protrusions for nine fossil hominid specimens. Given the fragile nature of the bones that make up the nasal cavity and how this diminishes the likelihood of their preservation in fossil crania, it was decided to take the least number of measurements needed to produce the prediction formulae. It would simply make no sense, in the context of ancient hominid facial approximation, to produce formulae that require measurements of intricate structures, such as those of the conchae of ethmoid, of extant species if these measurements could not be collected from fossils due to a severely low probability of preservation. Therefore, we only took measurements from aspects of the skull base and maxillofacial skeleton that are most durable and best protected against taphonomic deformation.

## Materials & Methods

The material used in this study consists of 19 computed tomography (CT) scans of chimpanzee (*P. troglodytes*) heads, previously analyzed in Campbell et al. (2021a), and 72 lateral cephalometric radiographs of humans. The chimpanzee sample was collected as Digital Imaging and Communications in Medicine (DICOM) format bitmap files from the Digital Morphology Museum, KUPRI (<u>dmm.pri.kyoto-u.ac.jp</u>). The sex ratio for the chimpanzee sample was 1:1.71 (7 male and 12 female) and the mean age 30.9 years (minimum = 9; maximum = 44; SD = 10.1). A complete list of all the chimpanzee subjects used in this study is presented in the S1 Table. The human sample was collected from the archive of a previous study (Simpson, 2005). The human sample includes two populations from different geographic areas: a Chinese population *(N* = 52), and an American/European population (*N* = 20). The sex ratio for the Chinese sample was 1:0.79 (29 male and 23 female). Exact ages were not available for this group but are classified as young adult. The sex ratio for the American/European sample was 1:0.82 (11 male and 9 female) and the mean age was 19 years and 1 month (minimum = 15; maximum = 32; SD = 4.7). Ethical approval was not required for the use of human subjects in this study due to the archival and anonymous nature of this material.

Measurements were taken on the midsagittal plane from the chimpanzee DICOM files in OsiriX MD, v. 11.02 (Visage Imaging GmbH, San Diego, USA), and from physical copies of the human radiographs with sliding and spreading calipers. Linear distances were collected using four standard cephalometric landmarks: basion (ba), nasion (n), pronasale (pn), and prosthion (pr) (Fig 1). These four landmarks were positioned onto the skulls and then measurements were taken for cranial base length (ba-n), nasal cavity length (ba-pn), and basion-prosthion length (ba-pr; hereafter referred to as jaw protrusion). All of these measurements were taken according to their descriptions in Martin and Saller (1957), which are listed in Table 1. Two angles were also measured (na-ba-pn and n-ba-pr) to examine the position of pronasale relative to the hard palate of the maxilla. The vertex of each triangle was positioned at basion with one ray to nasion. This was the starting point for both measurements. The second ray for each angle was positioned at pronasale for the first angle measurement and then to prosthion for the second angle measurement (Fig 1). Measurements were replicated seven days after initial assessment so that technical error or measurement (TEM), and relative TEM (rTEM) could be calculated from test-retest measurements. Measurement errors for all variables assessed in this study were less than 0.3% for chimpanzee CT scans and less than 0.9% for human radiographs.

**Fig 1.**
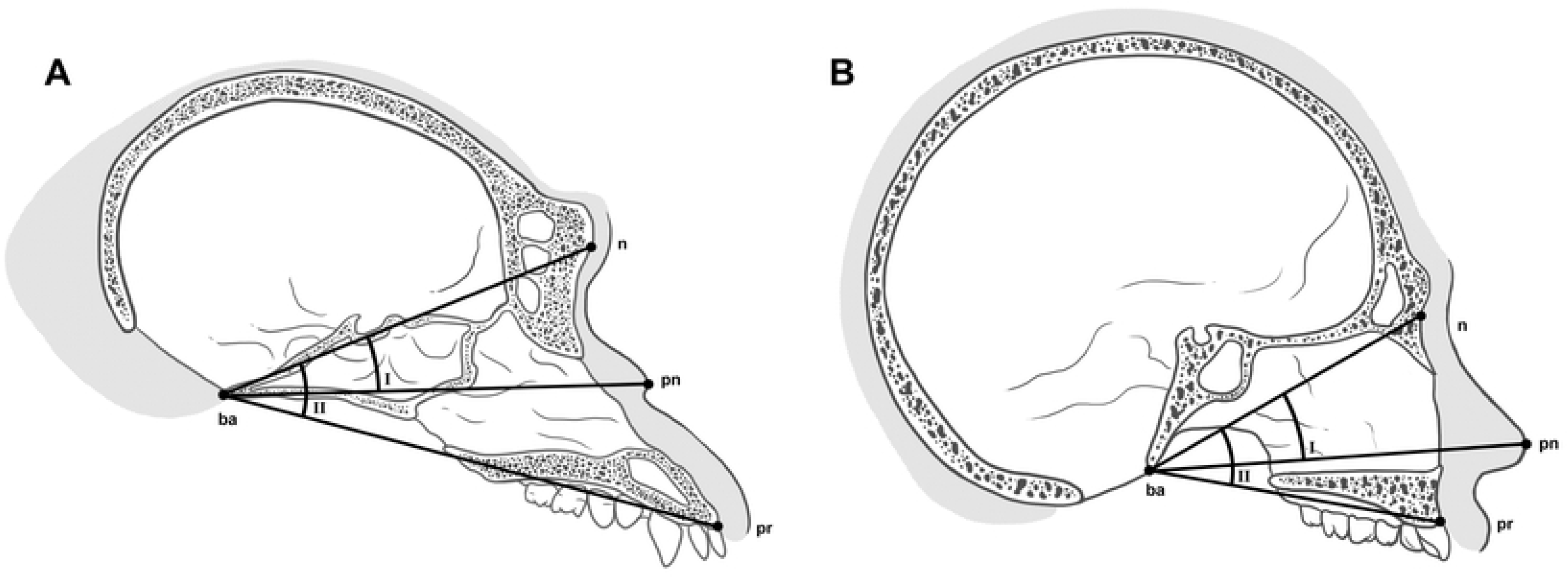
Locations of cephalometric landmarks used in this study and angles measured on the skull of a chimpanzee (*Pan troglodytes*; A) and modern human (*Homo sapiens*; B) in norma lateralis. (I) na-ba-pn angle. (II) na-ba-pr angle. See variable abbreviations in Table 1.

**Table 1.** Cephalometric landmarks used in this study including their abbreviations and definitions. Points are listed in alphabetical order for ease of reference.

| Landmark | Abbreviation | Definition |
| --- | --- | --- |
| **Basion** | ba | Anteriormost point of the foramen magnum in the midsagittal plane. |
| **Nasion** | n | Intersection of the nasofrontal suture in the median plane |
| **Pronasale** | pn | The most anterior point of the nose. |
| **Prosthion** | pr | The most anterior point of the maxilla in the midsagittal plane. |

Descriptive statistics were presented for all measurements and their ratios. We then used simple linear regression to examine the relationship of nasal size with cranial size. We regressed ba-pn against ba-n and assessed intra- and interspecific regression slopes and intercepts using 95% confidence intervals. We further regressed ba-pr against ba-n to compare this relationship with that of the previous for both species, as well as the na-ba-pn angle against the n-ba-pr angle. Reduced major axis (RMA) regression was used to produce the predictive equations because RMA, unlike ordinary least squared regression, is not influenced by random variation of individual measurements around the regression line (Sokal and Rohlf, 2012). All statistical analyses were carried out with the Statistical Package for the Social Sciences (SPSS®) software, v. 26.0 for Mac (SPSS Inc, Chicago, II, USA).

To demonstrate the reliability of the RMA prediction formulae, a set of out-of-group tests were performed. The first test was performed on two subjects that belonged to the same genus as the training sample, i.e., *Homo* (*n* = 1) and *Pan* (*n* = 1), and the second test, which functioned as an interspecies compatibility test, was performed on *Pan paniscus* (*n* = 1), *Gorilla gorilla* (*n* = 3), *Pongo pygmaeus* (*n* = 1), *Pongo abelli* (*n* = 1), *Symphalangus syndactylus* (*n* = 3), and *Papio hamadryas* (*n* = 3). Craniometric measurements were then collected from each specimen and used with the appropriate regression model to predict pronasale position. All subjects were collected from the Digital Morphology Museum, KUPRI (<u>dmm.pri.kyoto-u.ac.jp</u>), with the exception of the *Pan paniscus* subject, which was downloaded from Morphosource (https://www.morphosource.org), and the *Pongo abelli* subject. The *Pongo abelli* subject was scanned as part of a health assessment using the Siemens Biograph mCT PET/CT system at the South Australian Health and Medical Research Institute (SAHMRI). Slice thicknesses were set at 0.6mm and the animal was sedated and positioned in the supine position during the scanning procedure. The CT scans were then donated to the University of Adelaide for scientific research. A complete list of all out-of-group test material used in this study and their sources is presented in the S1 Table.

Exact ages for the infant *P. troglodytes* and *G. gorilla* are not provided by KUPRI, so we could only approximate their ages based on the dentition visible in their small immature jaws. In both *G. gorilla* and *P. troglodytes*, eruption of the first permanent molars occurs at approximately three years of age (Holly Smith et al., 1994). No permanent dentition eruption is visible in the *G. gorilla* subject, although the first permanent molars, canines, and incisors are approaching eruption. Therefore, PRI-7902 is not much less than 3 years of age. In the *P. troglodytes* subject, only the first permanent molars are fully erupted, so PRI-7895 is similarly approximated as 3 years of age.

In addition to the extant hominids, nasal protrusions were approximated for nine ancient hominid skulls using the RMA prediction formulae. Only hominid fossils with complete crania were selected. The crania included were two specimens representing the *Paranthropus* genus (KNM-WT 17000; *P. aethiopicus* and OH5; *P. boisei*), two specimens representing the *Australopithecus* genus, (Sts 5; *A. africanus* and MH1; *A sediba),* and five specimens representing the genus *Homo* (KNM-ER 1813; *H. habilis,* KNM-WT 15000; *H. ergaster* / *erectus,* LES1; *H. naledi,* Kabwe 1; *H. rhodesiensis* / *heidelbergensis,* and Amud 1; *H. neaderthalensis* / Neandertals). The soft tissue of each hominid was constructed in an oil-based modelling medium by GV using pegs anchored at basion to guide the shape of the nasal profiles and their underlying anatomy. A complete list of all fossil crania in this study and their sources is presented in the S1 Table.

## Results

Average cranial base length (ba-n) of chimpanzees is only slightly greater than that of modern humans (M = 107.86, SD = 5.83, n =19 and M = 104.28, SD = 5.71, n = 72 respectively), though T-test results show they formally differ significantly, *p* = 0.02 (2 tail). Average nasal cavity length (ba-pn) of chimpanzees (M = 131.49, SD = 8.03, n = 19) is somewhat greater than that of modern humans (M = 120.73, SD = 6.87, n = 72), *p* < 0.001 (2 tail), whereas average jaw protrusion (ba-pr) of chimpanzees (M = 149.62, SD = 11.20, n = 19) is much greater than that of modern humans (M = 98.14, SD = 6.11, n = 72), *p* < 0.001 (2 tail). Average na-ba-pr angle of chimpanzees (M = 35.23, SD = 3.46, n = 19) is only slightly smaller than that of modern humans (M = 41.72, SD 2.87, = 72), *p* < 0.001 (2 tail), as is average na-ba-pn angle of chimpanzees (M = 21.60, SD = 2.72, n = 19) compared to modern humans (M = 27.26, SD = 2.27, n = 72), *p* = 0.02 (2 tail).

The length of the nasal cavity (ba-pn) was, on average, equivalent to 122.1% and 115.8% of the length of the cranial base (ba-n) in chimpanzees and modern humans respectively. These rations were rather similar between species. In contrast, ratios of nasal cavity length (ba-pn) to jaw protrusion (ba-pr) were diametrically different. The ratio of mean jaw protrusion to the cranial base length was 138.7% in chimpanzees and 94.3% in modern humans. In other words, chimpanzees were observed to have a mouth that protrudes past the nasal cavity, whereas modern humans were found to have a nasal cavity that protrudes past the mouth even though lengths of the nasal cavity in both species are similar. These results and descriptive statistics for all measurements are illustrated in Fig 2 and Table 2 respectively.

**Fig 2.**
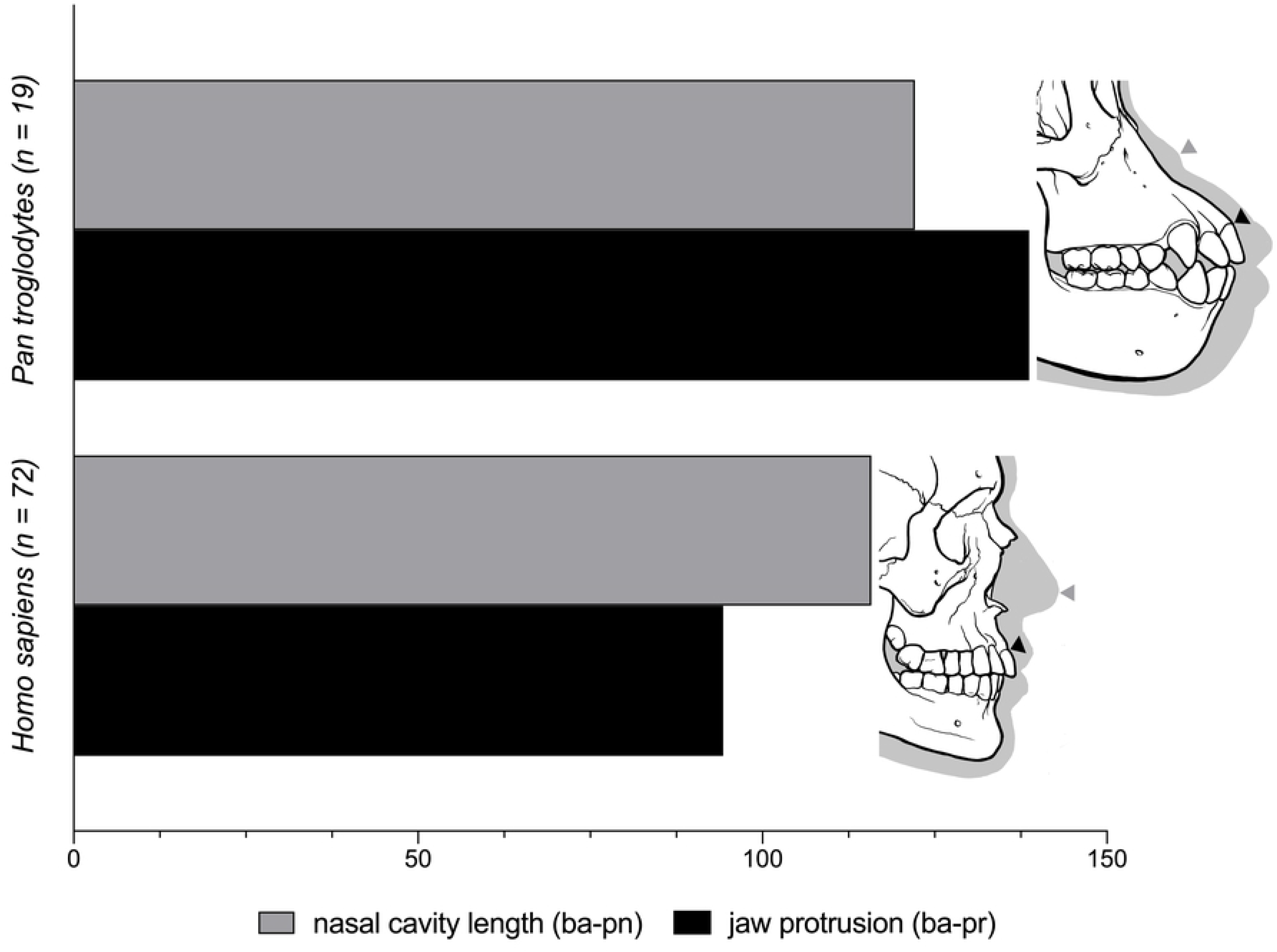
Comparison of nasal cavity length (ba-pn) to jaw protrusion (ba-pr) in chimpanzees (*Pan troglodytes*) and modern humans (*Homo sapiens*). See variable abbreviations in Table 1.

**Table 2.** Descriptive statistics for cranial base length (ba-n), nasal cavity length (ba-pn), and jaw protrusion (ba-pr) in mm for chimpanzees (*Pan troglodytes*) and modern humans (*Homo sapiens*). Angles measured in degrees are also shown.

| Variable<sup>a</sup> | Mean | SD | Minimum | Maximum |
| --- | --- | --- | --- | --- |
| **Chimpanzee (*n* = 19)** |  |  |  |  |
| ba-n | 107.86 | 5.83 | 98.00 | 123.00 |
| ba-pn | 131.49 | 8.03 | 115.90 | 143.10 |
| ba-pr | 149.62 | 11.20 | 122.90 | 163.10 |
| na-ba-pr angle | 35.23 | 3.46 | 30.26 | 43.99 |
| na-ba-pn angle | 21.60 | 2.75 | 16.54 | 26.92 |
| **Human (*n* = 72)** |  |  |  |  |
| ba-n | 104.14 | 5.62 | 92.80 | 119.20 |
| ba-pn | 120.57 | 7.07 | 106.30 | 140.80 |
| ba-pr | 98.14 | 6.11 | 85.30 | 114.40 |
| na-ba-pr angle | 41.72 | 2.87 | 35.50 | 48.00 |
| na-ba-pn angle | 27.26 | 2.27 | 21.70 | 32.00 |

<sup>a</sup> See variable abbreviations in Table 1.

Simple linear regressions revealed that nasal cavity length (ba-pn) was strongly and significantly correlated with cranial base length (ba-n) in both chimpanzees and modern humans (Table 3). In fact, the correlation coefficients obtained for each species were identical (*r* = 0.78; Table 3). Similarly, na-ba-pn and na-ba-pr angles were strongly correlated in both species with chimpanzees (*r* = 0.83) having a slightly greater correlation coefficient than that of modern humans (*r* = 0.73; (Table 3). Regression slopes were relatively consistent between species. Regressions of nasal cavity length (ba-pn) against cranial base length (ba-n) had a positive slope of 1.07 for chimpanzees and 0.94 for modern humans. Regressions of na-ba-pn angle against na-ba-pr angle had a positive slope of 0.65 for chimpanzees and 0.68 for modern humans. These results show that slopes are not species-specific, and neither are intercepts. Subsequent regression analyses combining *Homo* and *Pan* data clearly show this consistency across the entire sample with each combined regression providing a slightly better fit for both species (Fig 3).

**Table 3.** Ordinary least squares linear regressions of nasal cavity length (ba-pn) against cranial base length (ba-n) and na-ba-pr angle against na-ba-pn angle in chimpanzees and modern humans.

| Variable<sup>a</sup> | R | Slope | Intercept | CI<sup>b</sup><br>Slope<br>Lower | CI<br>Slope<br>Higher | CI<br>Intercept<br>Lower | CI<br>Intercept<br>Higher |
| --- | --- | --- | --- | --- | --- | --- | --- |
| **Chimpanzee (*n* = 19)** |  |  |  |  |  |  |  |
| ba-pn | 0.78* | 1.07 | 16.28 | 0.62 | 1.51 | -31.73 | 64.29 |
| na-ba-pn | 0.83 | 0.65 | -1.39 | 0.43 | 0.88 | -9.29 | 6.51 |
| **Human (*n* = 72)** |  |  |  |  |  |  |  |
| ba-pn | 0.78* | 0.94 | 22.89 | 0.76 | 1.12 | 4.12 | 41.66 |
| na-ba-pn | 0.73 | 0.68 | -1.10 | 0.58 | 0.78 | -5.14 | 2.93 |

<sup>a</sup> See variable abbreviations in Table 1.
<sup>b</sup> 95% confidence interval.
\* Indicates where correlations coefficients were identical between species.

**Figure 3.**
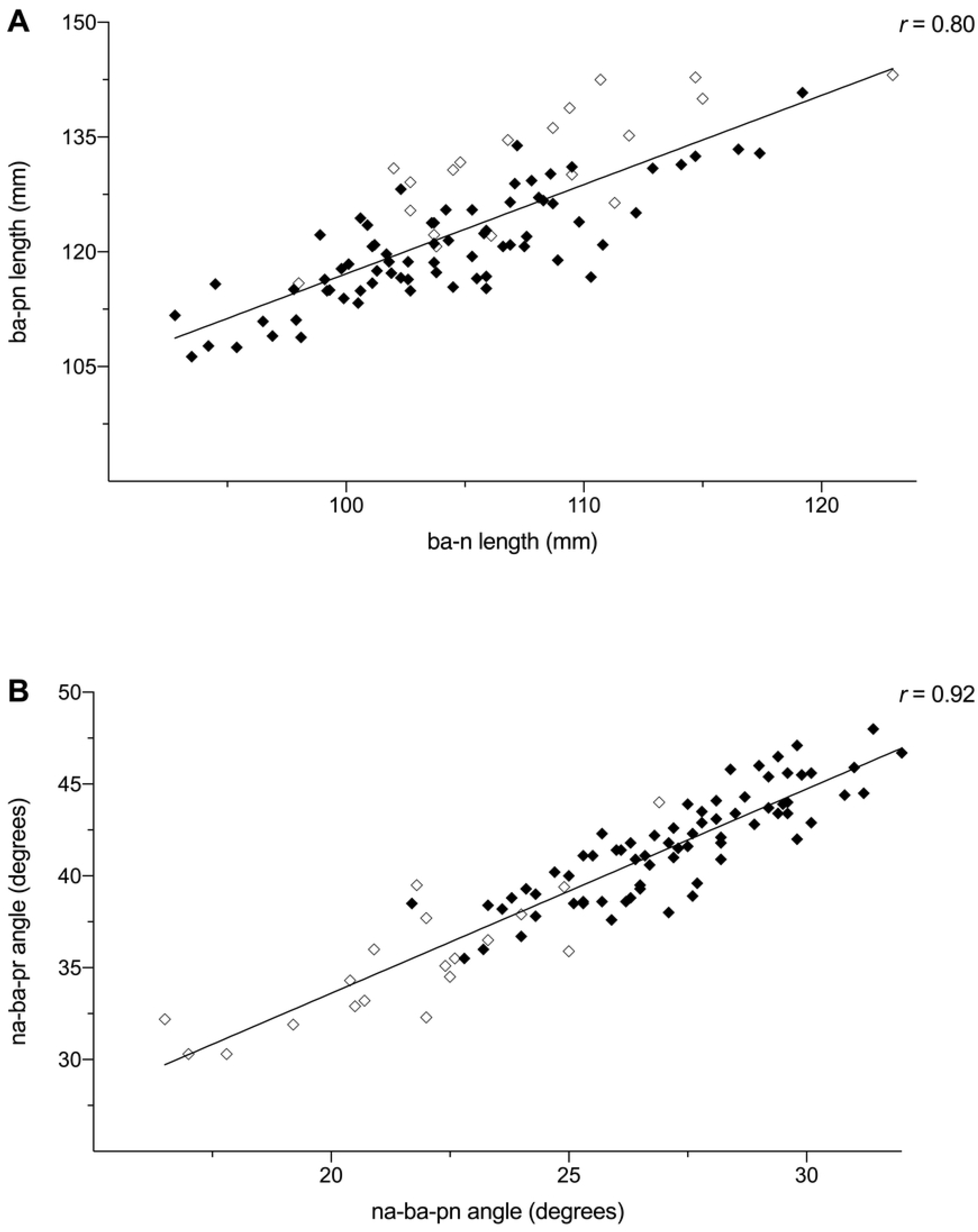
Bivariate scatterplots showing regressions for a combined sample of chimpanzees (*Pan troglodytes*; ◊) and modern humans (*Homo sapiens*; ◆). (A) Regression of nasal cavity length (ba-pn) on cranial base length (ba-n). (B) Regression of na-ba-pr angle on na-ba-pn angle. See variable abbreviations in Table 1.

Given that simple linear regressions were able to identify statistically significant correlations in the combined sample, as well as establish homogeneity of interspecific covariation, we transformed the prediction equations using Reduced Major Axis (RMA) regressions to remove the influence of individual variation on predictions. Nasal protrusions for modern humans and chimpanzees could thus be approximated using the following equations:

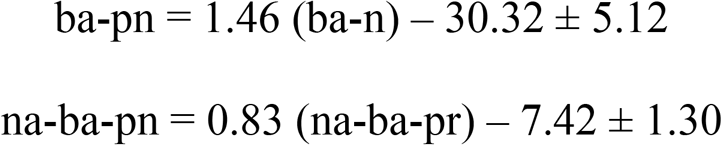

The results of the out-of-group test using the above RMA regression formulae on one member of *Homo sapiens*, and one member of *Pan troglodytes* were quite accurate. The average difference between actual and predicted ba-pn length and na-ba-pn angle for the human subject was 1.9 mm and 1.4 mm for the chimpanzee subject. The interspecies test results performed on *Pan paniscus*, *Gorilla gorilla*, *Pongo pygmaeus*, *Pongo abelli*, *Symphalangus syndactylus*, and *Papio hamadryas* using the same formulae are shown in Fig 4. We found a substantial difference in the predictive accuracy of the regression models among species. The performance of the models produced accurate results for *P. paniscus* (*n* = 1) and *G. gorilla* (n = 3), but poor results for *P. pygmaeus* (*n* = 1), *P. abelii* (*n* = 1), *S. syndactylus* (*n* = 3) and *P. hamadryas* (*n* = 3). This clearly indicates that there is an incremental decline in the predictive accuracy of the equations depending on the phylogenetic distance of these species relative to *H. sapiens* and *P. troglodytes*.

**Fig 4.**
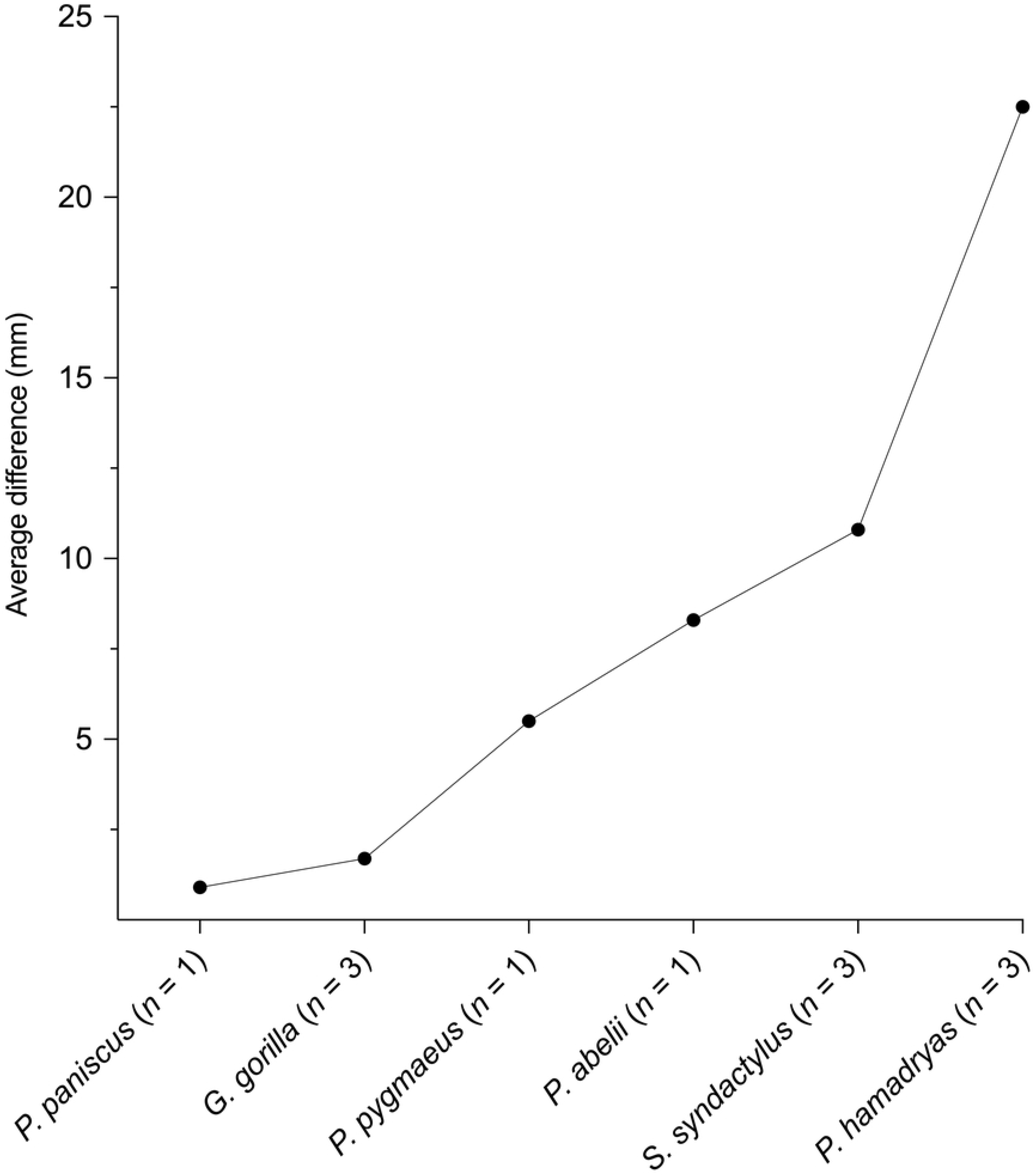
Average difference between predicted and ground truth values shown for the out-of-group tests performed on six separate species that are outside of the chimpanzee/human training sample. Notice the influence of the phylogenetic position of each species relative to modern humans and chimpanzees, i.e., from Hominoidea to Cercopithecoidea, and how this leads to a progressive increase in approximation error.

It is important to emphasise the ages of the out-of-group test subject. The age of the infant *Pan troglodytes* subject was 3 years, the *P. paniscus* subject 4 years, the infant *G. gorilla* 3 years, the male *G. gorilla* 46 years, and the female *G. gorilla* 54 years. All these subjects were therefore outside of the age-range of the chimpanzee/human training sample. Given that the regression formulae were able to provide quite accurate estimates for all these subjects, it appears that when the regression formulae are compatible with a given species, they do not appear to be restricted to a specific age-range.

The results of the regression formulae applied in 3D approximations of the nasal region for members of *H. sapiens*, *P. troglodytes*, *P. paniscus*, and *G. gorilla* are shown in Fig 5. The formulae of the present study have allowed for the objective and accurate approximation of the soft tissue landmark pronasale for all these species from measurements of their bone alone. 3D approximations were not performed for *P. pygmaeus*, *P. abelii*, *S. syndactylus*, and *P. hamadryas* because, as stated above, the prediction formulae produced poor estimates and are therefore incompatible with these species.

**Fig 5.**
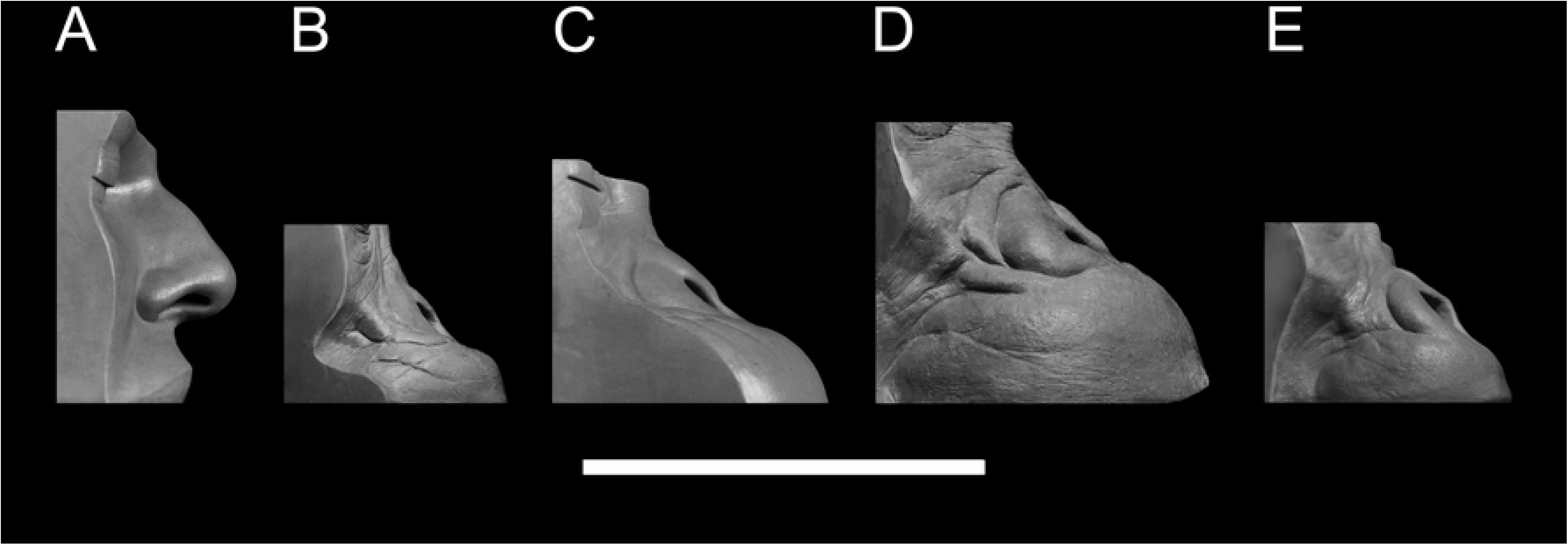
Reduced major axis regression formulae applied in 3D approximations of the nasal region for out-of-group test subjects in norma lateralis. (A) *H. sapiens*: Anonymous 29-year-old male subject. (B) *P. troglodytes*: PRI-7895, 3-years-old. (C) *P. paniscus*: S9655, 4-years-old. (D) *G. gorilla*: PRI-Oki, 54-years-old. (E) *G. gorilla*: PRI-7902, 3-years-old. Scale Bar = 10 cm.

The results of the regression formulae applied in 3D approximations of the nasal regions for extinct hominid are shown in Fig 6. The results are consistent with previous interpretations of these species (Franciscus and Trinkaus, 1988b; Gurche, 2013). There is significant variation in the nasal profile among hominid clades, from the chimp-like profiles of Pliocene *Australopithecus* to the human-like profiles of Pleistocene Neandertals. Since we have observed that regression formulae derived from modern human and chimpanzee material demonstrate high reliability when applied to all African great apes, i.e., bonobos and gorillas, we put forward the hypothesis that the same formulae are applicable to ancient Plio-Pleistocene hominids and, since the approximations were not produced using artistic intuition, that our results are empirically and scientifically accurate.

**Fig 6.**
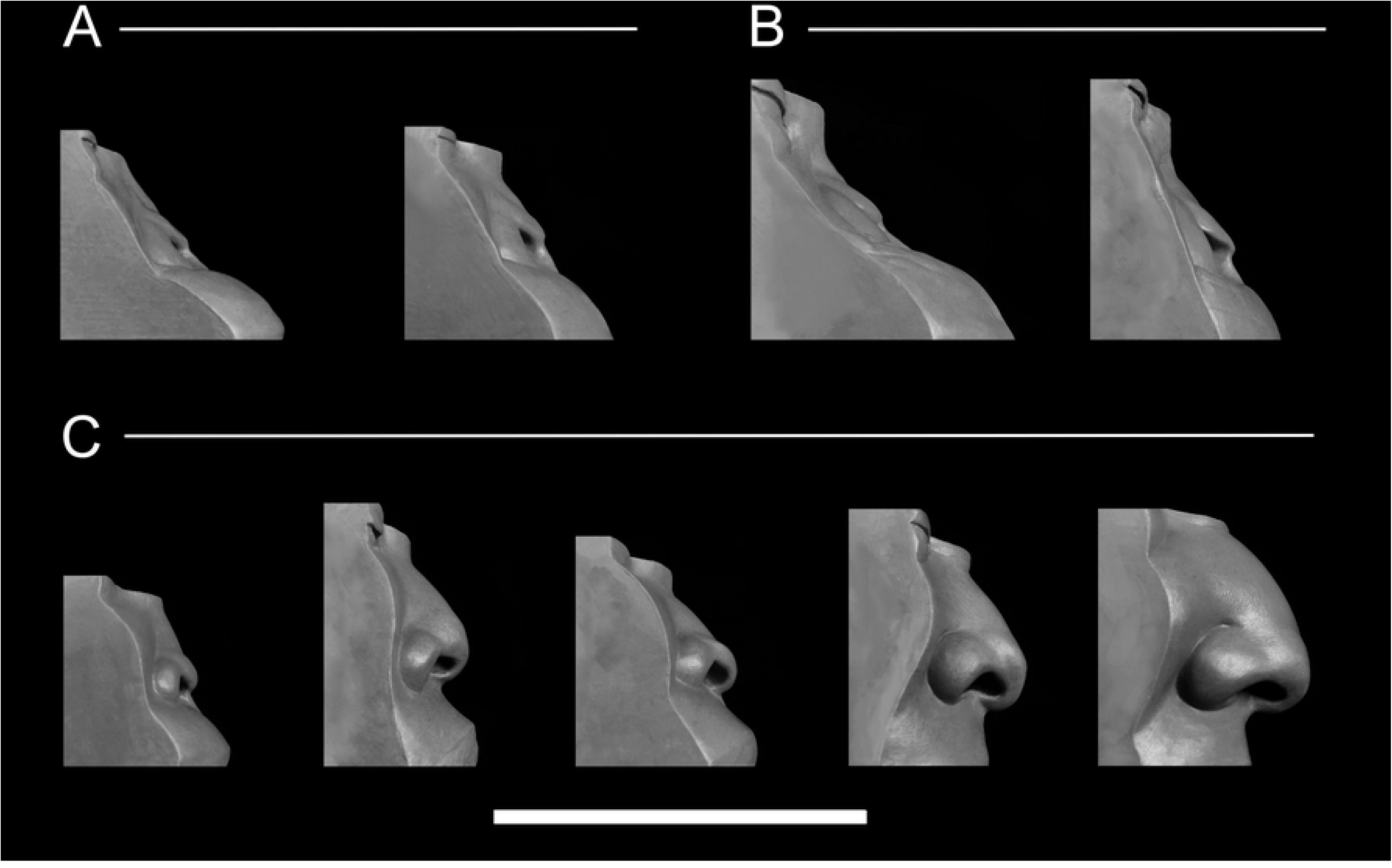
Reduced major axis regression formulae applied in 3D approximations of the nasal region for extinct hominids in norma lateralis. (A) *Australopithecus* genus: Sts 5 (*A. africanus*) and MH1 (*A. sediba*). (B) *Paranthropus* genus: KNM-WT 17000 (*P. aethiopicus*) and OH5 (*P. boisei*). (C) *Homo* genus: KNM-ER 1813 (*H. habilis*), KNM-WT 15000, (*H. ergaster* / *erectus*), LES1 (*H. naledi*), Kabwe 1 (*H. rhodesiensis* / *heidelbergensis*), and Amud 1 (*H. neaderthalensis* / Neandertals). Scale Bar = 10 cm.

## Discussion

Recently, Campbell et al. (2021a) showed that facial soft tissue thicknesses covary with craniometric dimensions in chimpanzees. However, Campbell et al. (2021a) only analyzed chimpanzees and their formulae provide no information about the nasal protrusions of archaic hominids. The present study begins to alleviate this, at least in part. We have identified another set of predictable relationships, only this time for predicting pronasale position. Given that chimpanzee and modern human means differ significantly, it is not possible to approximate the position of pronasale for these species using averages. More and more forensic studies are revealing that linear regression models actually outperform averages (Campbell et al., 2021a; Dinh et al., 2011) and this study suggests no different.

Focusing on the interspecies out-of-group tests, taking the phylogenetic position of each species into account provides a sound explanation for the decrease in performance observed in each regression model. As can be seen in Fig 7, the regression trajectories of *P. paniscus* and *G. gorilla* appear to share a very close affinity with the chimpanzee/human training sample, whereas *P. pygmaeus, Pongo abelli, S. syndactylus*, and *P. hamadryas* appear to have group-specific slopes and intercepts. What is most surprising about these results is the *G. gorilla* sample. The RMA formulae predictions for the infant and two adult subjects produced accurate results, which not only highlights the validity of correlations identified in the present study but also, and more importantly, their compatibility with individuals of different ages belonging to separate species.

**Fig 7.**
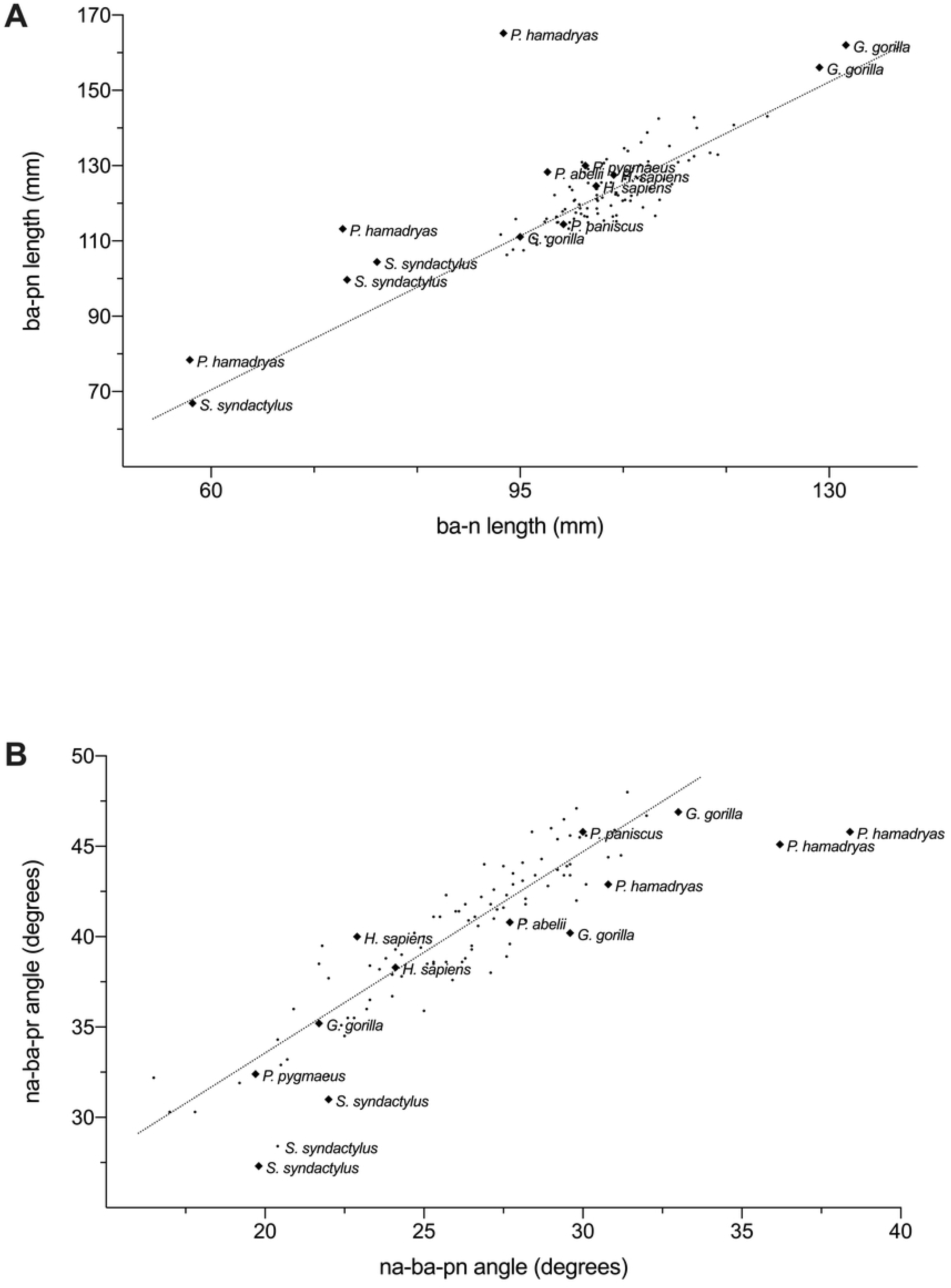
Bivariate scatterplots with actual values for pronasale position in *Pan paniscus* (*n* = 1), *Gorilla gorilla* (*n* = 3), *Pongo pygmaeus* (*n* = 1), *Pongo abelli* (*n* = 1), *Symphalangus syndactylus* (*n* = 3), and *Papio hamadryas* (*n* = 3) superimposed over the chimpanzee/modern human regression lines. (A) Regression of nasal cavity length (ba-pn) on cranial base length (ba-n). (B) Regression of na-ba-pr angle on na-ba-pn angle. See variable abbreviations in Table 1.

Based on the results of the out-of-group interspecies compatibility tests, we suggest that it can be formulated as a general rule that hominids with longer cranial base lengths tend to have longer nasal cavities, and that hominids with maxillae tilted further down from the Frankfurt horizontal plane tend to have axes of the nasal cavity also directed further downwards. Furthermore, since this has been identified in two extant species of hominid, which feature quite distinct skull morphologies, and that the regression models can reliably approximate pronasale position and nasal protrusion in other African great apes (i.e. *P. paniscus* and *G. gorilla*), these equations can be applied in facial approximation of extinct Plio-Pleistocene hominins. Furthermore, given the absence of soft tissue in the fossil record, which has been shown to pose a particular problem for approximations of *Homo habilis* and *Homo naledi* (Campbell et al., 2021b), quantitative linear regression essentially removes descriptive speculation during the approximation of this aspect of the nose for these species.

Owing to the relatively similar measurements of nasal cavity length among the individuals in our chimpanzee and modern human sample, our results are congruent with Nishimura et al. (2016) in that that projecting noses are only partially a result of local adaptations to climate in the genus *Homo* (Franciscus and Trinkaus, 1988a). To elaborate on this, consider that during human evolution there was a change in habitat from tropical rainforest to a greater variety of possible habitats, such as more open savannah and woodland mosaics (Lovejoy, 1981). Chimpanzees, on the other hand, provide a case of evolutionary stasis (Diogo et al., 2017), in large part due to not migrating out of their ancestral environments. Despite this, nasal cavity lengths are similar between chimpanzee and modern human species and thus one cannot accept the hypothesis that local adaptations to climate included nasal cavity lengths in our sample. It seems that nasal cavity size has been retained without change in hominin evolution while reductions of the masticatory apparatus during human evolution revealed the prominent nose exhibited in modern humans. This is unlike all other non-human apes who lack a prominent nose because they reach dimensions of jaw protrusion far exceeding those found in modern humans. Unlike modern humans, chimpanzees did not evolve the same repertoire of extra-oral methods for predigesting food and, therefore, did not undergo any reductions in their masticatory apparatus. Furthermore, their social relationships based on male dominance did not allow canine reduction and loss of the C/P3 honing complex (Delezene, 2015). In contrast to changes in food preparation and canine use in competition for dominance, neither of the species evolved any extra-nasal methods for conditioning inspired air, which is the most likely explanation for why dimensions of nasal cavity are so strikingly similar between modern humans and chimpanzees. Numerous clinically oriented studies have shown that there are at least two functions of the nose; humidification and temperature modification of inspired air (Ewert, 1965; Walker and Wells, 1961). Nasal cavity length was clearly formed by natural selection for these adaptive roles, but the fact that nasal cavity lengths between humans and chimpanzees are so similar shows that these changes were minimal relative to changes in the size of the masticatory apparatus

Our analyses also concur with the observation that modern human facial growth is retarded relative to chimpanzees (Johanson, 1981; Penin et al., 2002). This observation has been explored elsewhere in the neotenic theory of the human skull (Gould, 1977), and the self-domestication hypothesis (Theofanopoulou et al., 2017). We will not go into detail about these subjects here, but it suffices to say that during chimpanzee, as well as other great ape, ontogeny, the degree of facial prognathism increases from infancy to adulthood. In contrast, modern human skulls appear paedomorphic relative to chimpanzees; this shared affinity between adult humans and sub-adult chimpanzees is only temporary since great apes do not retain this morphology into adulthood (Johanson, 1981). This is to say, that the developmental changes that occur throughout chimpanzee ontogeny, which result in a mouth that protrudes past the nose, do not occur in modern humans. It is thus reasonable to assume that if facial prognathism would have persisted in the genus *Homo*, the noses of modern humans today would also appear flat like those of chimpanzees.

Our approximations of Plio-Pleistocene hominids shown in Fig 6 favor the hypothesis that the length of the nasal cavity remained relatively constant throughout human evolution. The unique projecting nose exhibited in anatomically modern humans appears not to have been the result of the cartilaginous components of the nose actively growing out of the face as an environmental adaptation. Instead, our approximations suggest that projecting noses were simply the result of reductions in the masticatory apparatus over time. This point is made obvious by comparing the position of pronasale relative to prosthion between approximations in Fig 6. The more superior position of pronasale and anterior position of prosthion in relation to the piriform aperture, the more chimp-like the nose appears. In direct contrast, the more inferior position of pronasale and superior position of prosthion, the more human-like the nose appears. Interestingly, two hominids (KNM-ER 1813; *Homo habilis* and LES1; *Homo naledi*) are not so easily classified as they appear to represent a morphology that is neither ape-like nor human-like. However, the approximations are not outside the realm of possibility of what could have been present in the morphology of intermediate species.

A limitation of this study is the low number of individuals in the chimpanzee sample and out-of-group tests. This is not uncommon for studies of great ape soft tissue due to their sparse availability. It is an unfortunate predicament that researchers find themselves in when studying the soft tissue parts of these endangered and protected animals. Osteological material is plentiful but soft tissue is relatively non-existent. As invaluable as the primate data from KUPRI are, there exists a need for the expansion of publicly available data to include a larger number of living individuals. One possible solution to this problem is to collect existing data from primate sanctuaries and zoos since it is reasonable to speculate that a large number of these animals would have been scanned over the years for various health reasons. If these data could be collected and made freely available, it would not only benefit researchers but also the public in providing an unprecedented look into the anatomy of our closest living relatives. The Visible Ape Project is a recently developed resource that aims to accomplish this feat (Barger et al., 2021), although partnerships with veterinarians and conservationists are proving difficult due to the personal attachment of keepers to these animals and related hesitancy sharing these data. Furthermore, the duplication and scattering of data into many separate online repositories may result in a persistent struggle to keep these data orderly and to track their primary sources.

The other limitation of this study is that it only analyzed the length of the nose, which excludes other elements of the nasal form lateral to the mid-sagittal plane. Although pronasale position is the most defining point for the lateral view of the nose, other features of the nose are arguably as important as the protrusion alone. Alar and nostril size and shape have been somewhat investigated in modern humans but there is still a very wide-open field for researchers to describe these characters in non-human apes. To the knowledge of the authors, Hofer (1972) is the only one to have formally studied and published descriptions of the soft tissue parts of the nose in nonhuman apes. In his analysis of gorilla, Hofer (1972) recorded many interesting observations found in their noses, including a number of sulci of the oro-nasal region that are not present in modern humans. The causes for these differences are unclear, so it is worth investigating this material further as explanations for these variations may provide clues to the possible presence of these features in ancient hominids. As such, this points to a large void in the literature requiring that other aspects of the nose receive equal attention in future studies.

## Conclusions

Facial approximations of Plio-Pleistocene hominids provide a fascinating insight into the possible appearance of our ancestors, which helps to encourage interest in human evolution. However, there is considerable variability in the appearance of current approximations of the same individual. This is in large part due to unreliable approximation methods, especially concerning the facial features such as the nose. The main conclusions that can be drawn from this study is that there are homogenous relationships between the skull and the soft tissue parts of the nose in both chimpanzee and modern human species. Regressions combining chimpanzee and modern human measurements have shown that this amalgam of data can produce statistically reliable prediction formulae. These prediction formulae have some degree of interspecies compatibility, since we have shown that the formulae can be applied to chimpanzees, modern humans, bonobos, and gorillas. Based on these results, we hypothesize that the same formulae are valid for approximating the nasal profiles for extinct hominids because their craniums fit into the range of variability present in our sample of extant hominid species. However, our research has limitations in that it produced regression models using chimpanzee and modern human material, and only analyzed the protrusion of the nose. More investigations are therefore needed to rediscover these relationships in larger samples of great apes, such as orangutans, and to produce prediction formulae for other measurements of the soft parts of the nose, such as the ala nasi. Nevertheless, the present comparison of chimpanzee and human nasal cavity lengths shows that these species are more similar to each other than they are different and that the regression formulae can be safely applied to all African apes. Lastly, our study does not support the view that the nose arose out of the face. Rather, it provides evidence supporting the alternative hypothesis that it was reductions in the size of the masticatory apparatus over time that led to the external appearance of the anterior part of the nasal cavity in the genus *Homo*.

## Supporting information

S1 Table. List of non-human primate subjects included in this study.

(DOCX)

## Acknowledgments

We would like to thank the Kyoto University Primate Research Institute (Kyoto, Japan) and Dr Ellie Simpson for facilitating the acquisition of the data used in this study. We would also like to thank Victor Surovec and Daniel Collins of Arizona State University for providing access to the Makerspace and the Vizproto Lab during the production of the approximations presented in Figs 5 and 6. We would like to acknowledge David McLelland from Zoos South Australia and Georgia Williams for facilitating the acquisition of Puspa’s CT scan. The authors acknowledge the facilities, scientific and technical assistance of the National Imaging Facility, a National Collaborative Research Infrastructure Strategy (NCRIS) capability, at the Large Animal Research and Imaging Facility, South Australian Health and Medical Research Institute. We would also like to thank John Engelhardt for the illustration presented in Fig 1.

